# Activity-dependent nucleation of dynamic microtubules at presynaptic boutons is required for neurotransmission

**DOI:** 10.1101/679464

**Authors:** Xiaoyi Qu, Atul Kumar, Heike Blockus, Clarissa Waites, Francesca Bartolini

## Abstract

Control of microtubule (MT) dynamics is critical for neuronal function. Whether MT nucleation is regulated at presynaptic boutons and influences overall presynaptic activity remains unknown. By visualizing MT dynamics at individual excitatory *en passant* boutons in axons of hippocampal neurons we found that MTs preferentially grow from presynaptic boutons as a result of γ-tubulin and augmin-dependent nucleation. MT nucleation at boutons is promoted by neuronal activity, functionally coupled to synaptic vesicle (SV) transport, and required for neurotransmission. Hence, *en passant* boutons act as hotspots for activity-dependent MT nucleation, which is required for neurotransmission by providing the tracks for a rate-limiting supply of SVs to sites of neurotransmitter release.

**Highlights:**

- Excitatory boutons are hotspots for neuronal activity-induced γ-tubulin dependent MT nucleation
- The augmin complex is required for the correct polarity of presynaptic *de novo* nucleated MTs
- Presynaptic MT nucleation promotes SV motility and exocytosis at sites of release

**In Brief:** Our results demonstrate that excitatory *en passant* boutons are hotspots for neuronal activity-induced γ-tubulin- and augmin-dependent oriented MT nucleation, and that the resulting presynaptic *de novo* nucleated MTs promote inter-bouton SV motility which is rate-limiting for neurotransmitter release.

## INTRODUCTION

Dynamic microtubules (MTs) play a crucial role for rapid forms of axonal and dendritic transport and synaptic transmission (*1–3*). At postsynaptic sites, MT invasion into spines regulate spine morphology, motor/cargo pair transport into spines, and synaptic activity (*4–9*).

MT dynamics and organization at presynaptic boutons, however, remains a largely uncharted area due to the limited size of the boutons in most mammalian neurons and loss of dynamic MTs using standard fixation procedures. Classic electron microscopy studies have described the presence of pools of MTs at presynaptic sites in the rat cortex and cerebellum, including synaptic vesicle (SV)-clothed MTs attached to the active zone and peripheral coils of MTs in close contact with presynaptic mitochondria (*10–16*). These observations suggested that presynaptic MTs may be functionally coupled with synaptic transmission by: (1) driving the interbouton translocation of SVs (2) maintaining the organization of the presynaptic active zone (AZ), (3) contributing to the maturation of postsynaptic receptors by conveying new SVs to be released at certain presynaptic sites (*17*), (4) anchoring mitochondria at sites of release (*14, 18*). Few studies, however, have explored these possibilities using modern imaging approaches. Presynaptic MAP-associated stable MT loops, EB1-labeled dynamic pioneer MTs, and DAAM-dependent stable MT regulation of SV release and AZ morphology have been described at the *Drosophila* neuromuscular junction (*19–21*). In addition, a marginal band of modified stable MTs residing in the giant synaptic terminal of the goldfish retinal bipolar neuron mediates mitochondria transport and organization at synaptic terminals (*18*). Finally, presynaptic MTs appear to regulate interswelling SV transport in mouse giant calyceal terminals (*22*), and interbouton dynamic MTs have been recently implicated in Kif1A-dependent delivery of SVs in hippocampal neurons (*23*). It remains unknown, however, whether axonal *de novo* microtubule nucleation occurs post-development, is required for neurotransmission and a feature of neurons residing in an intact circuit.

Here we describe that dynamic MTs emerge at excitatory presynaptic *en passant* boutons as a result of restricted γ-tubulin- and augmin-dependent *de novo* MT nucleation, with γ-tubulin regulating the nucleation density, and augmin directing the uniform, plus-end directed growth towards the distal end of the axon. Moreover, we found that *de novo* nucleation of dynamic MTs at boutons is conserved in an intact circuit, stimulated by neuronal activity and required for neurotransmission by providing the tracks for interbouton transport of a rate-limiting pool of SVs at sites of neurotransmitter release.

## RESULTS

### Dynamic MT plus ends emerge from presynaptic boutons in axons of hippocampal neurons *in vitro* and *ex vivo*

We examined the dynamic behavior of presynaptic MTs in axons of pyramidal neurons by transfecting cultured hippocampal neurons (18DIV) with EB3-GFP and vGlut1-Cherry or VAMP2-Cherry constructs. Fast time-lapse imaging of EB3 labeled MT plus ends allows the quantitative assessment of MT dynamics at or away from individual glutamate release sites labeled with vGlut1-mCherry (Figure 1A and Figure S1A). Presynaptic dynamic MTs were distinguished based on their plus end contact with vGlut1^+^ boutons and defined as: 1) *interbouton* MTs that have no contact with the presynaptic markers, or 2) *intrabouton* MTs that interact with the presynaptic synaptic vesicle marker at any point during their lifetime. Intrabouton MTs were further classified as: (1) nucleating/rescuing (starting) at boutons, (2) undergoing catastrophe/pausing (ending) at the boutons or (3) passing through the bouton (Figure 1A). Based on kymograph analyses of EB3 comet motion relative to the stable pool of two distinct presynaptic markers (vGlut1 and VAMP2; Figure S1A), we found a higher density of longer lived intrabouton MTs than interbouton MTs in untreated neurons (Figure 1B and Figure S1A,B). When we further analyzed the pool of intrabouton MTs that end or start at a bouton by measuring their observed frequencies compared to their relative probability predicted by chance, we found a significantly higher frequency of EB3 comets starting rather than ending at presynaptic boutons (Fig. 1C). This suggests that MT nucleation labeled by EB3 comets are frequently initiated at presynaptic boutons. To examine if this result is also observed *in situ* at synapses in a native circuit, we performed comparable time-lapse imaging of EB3-labeled dynamic MTs in close contact with stable vGlut1-labeled excitatory presynaptic boutons in the CA1 region of acute hippocampal slices isolated from 21 day old mice that had been *in utero* co-electroporated with EB3-EGFP and vGlut1-mCherry at E15.5 (Figure 1D,E and Video S1). We observed a similar level of increase in frequency of EB3 comets starting but not ending at boutons compared to predicted random probability (Figure 1F), confirming the preferential regulation of EB3 comets initiating at presynaptic boutons *ex vivo* where cellular architecture and synaptic connectivity is largely intact.

**Figure 1.**
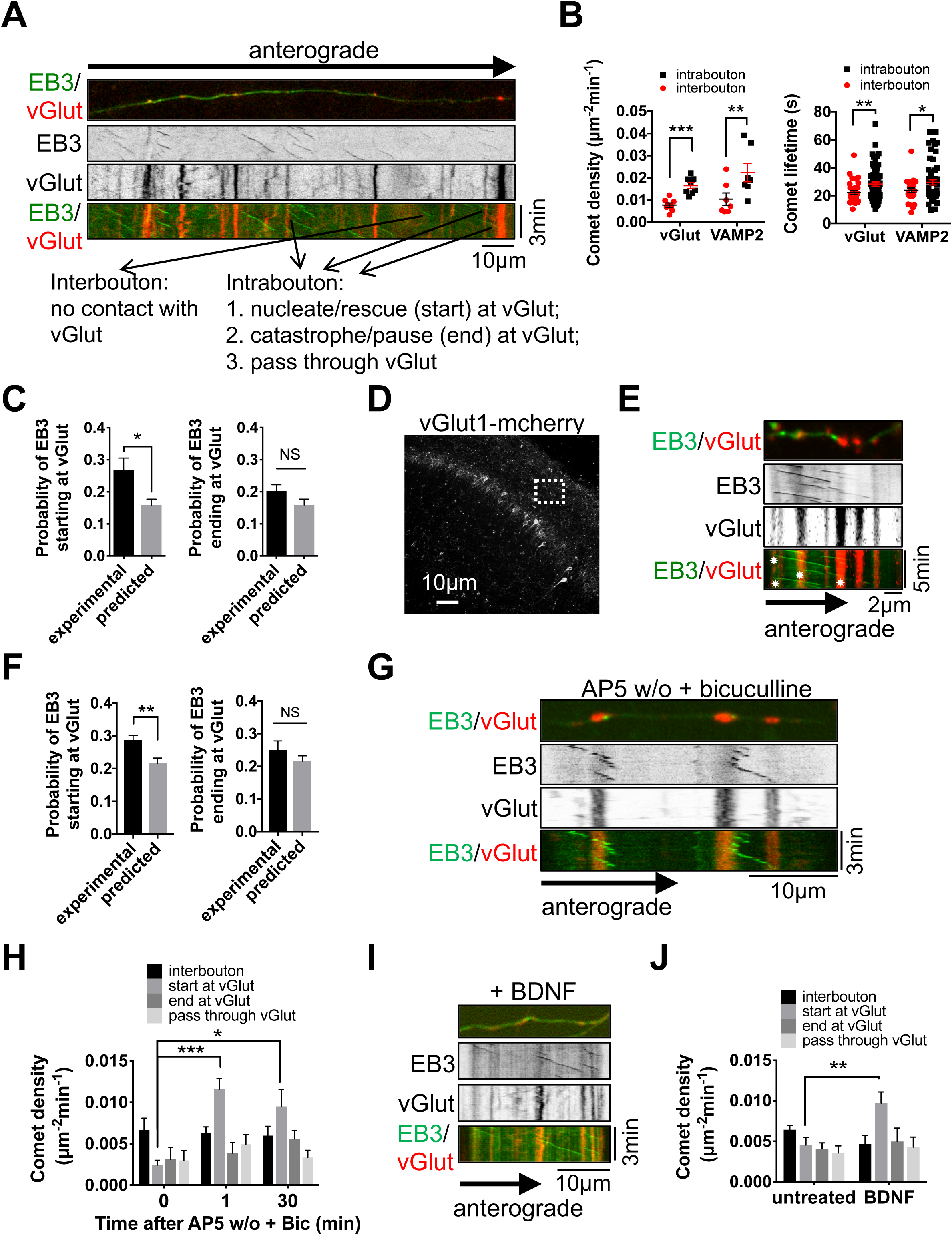
Dynamic MT plus ends initiate at presynaptic boutons upon induction of neuronal activity. (A) Representative maximum projection of spinning disk confocal fluorescence images and kymographs of untreated hippocampal neurons (21DIV) transfected with EB3-EGFP (EB3) and vGlut1-mCherry (vGlut) 24h prior to live imaging. Shown by arrows are the classifications of axonal dynamic MTs in inter- and intrabouton MTs as described. (B) Comet density and lifetime of intrabouton and interbouton EB3 comets in neurons treated as in (A). (C) Probability of EB3 comets starting or ending at vGlut^+^ stable puncta from experimental observations or predicted random events based on observed average density of vGlut^+^ stable puncta per axonal length (µm). Predicted probability refers to the chance of any EB3 comet to randomly start or end at vGlut, assuming that the diameter of a vGlut^+^ stable punctum is 1µm. (D) Representative maximum projection of spinning disk confocal fluorescence images of acute hippocampal slice in CA1 region from 21 day old mice electroporated with EB3-EGFP and vGlut1-mCherry at E15.5. vGlut1-mCherry channel under a 20x objective is shown and the white dotted box indicates the axonal region selected for live imaging using a 60x objective. (E) Representative maximum projection of spinning disk confocal fluorescence image and kymographs of an axon from boxed region in D. Asterisks indicate comet tracks initiating from vGlut^+^ stable boutons. (F) Probability of EB3 comets starting or ending at vGlut^+^ stable puncta from experimental observations or predicted random events in axons from acute hippocampal slices. (G,I) Representative kymographs of hippocampal neurons (21DIV) transfected with EB3 and vGlut and pretreated with 50µM of the NMDA receptor antagonist D-AP5 6-12h prior to imaging, followed by a 1min washout (w/o) and incubation with 20µM of the GABAa receptor antagonist bicuculline up to 30min (G) (Figure S1C), or directly treated with 50ng/mL BDNF for 1min (I) (Figure S1G). (H, J) Subclassified comet density for intrabouton MTs measured in hippocampal neurons treated as in G or I for the indicated times. * p<0.05; ** p<0.01; *** p<0.001 by two-tailed Student’s t-test (B, C: N = 5-8 axons; F: N = 5 axons; H, J: N = 6-9 axons). NS, non significant.

We tested whether initiation of EB3 comets at boutons was regulated by action potential (AP)-driven neuronal activity, and measured EB3 comet density starting at vGlut1^+^ boutons upon either acute AP5/bicuculline treatment to increase AP firing (Figure 1G,H, Figure S1D, and Video S2) or BDNF-induced neuronal activation (Figure 1I,J, Figure S1F, and Video S3). Strikingly, both treatments resulted in a significant and specific increase in the density of comets initiating at vGlut1^+^ boutons starting after 1min of treatment, but not in D-AP5 washout or untreated control conditions (Figure S1C). In addition, the percentage of nucleated comets reaching the next distal bouton increased by 3-fold upon acute neuronal firing (Figure S1E). Together, these data demonstrate that initiation of dynamic MT plus ends preferentially occurs at presynaptic sites, shows biased directionality being almost always oriented towards the distal tip of the axon, and can be induced by neuronal activity.

### Activity-promoted initiation of distally oriented dynamic MT plus ends at boutons results from γ-tubulin and augmin-dependent *de novo* MT nucleation

The increase in distally oriented EB3 comets could either result from *de novo* nucleation by γ-tubulin or from polymerization from preexisting MTs by the activity of MT rescue factors. In axons of developing neurons, γ-tubulin is required for acentrosomal MT organization and the augmin complex interacts with γTuRC to control MT distal polarity in axons (*24, 25*). However, whether axonal MT nucleation in the shaft and/or at presynaptic contacts occurs in axons of neurons with mature synapses is currently unknown. First, we examined endogenous γ-tubulin localization relative to the presynaptic marker synapsin1 in 21DIV hippocampal neurons, and found that ∼80% of γ-tubulin signal colocalized with synapsin1^+^ puncta compared to only ∼20% of GAPDH cytosolic control (Figure 2A,B). Similarly, ∼80% of exogenously expressed γ-tubulin-emerald co-localized with stable vGlut1-Cherry^+^ (vGlut) puncta in 21DIV hippocampal neurons (Figure S2A,B).

**Figure 2.**
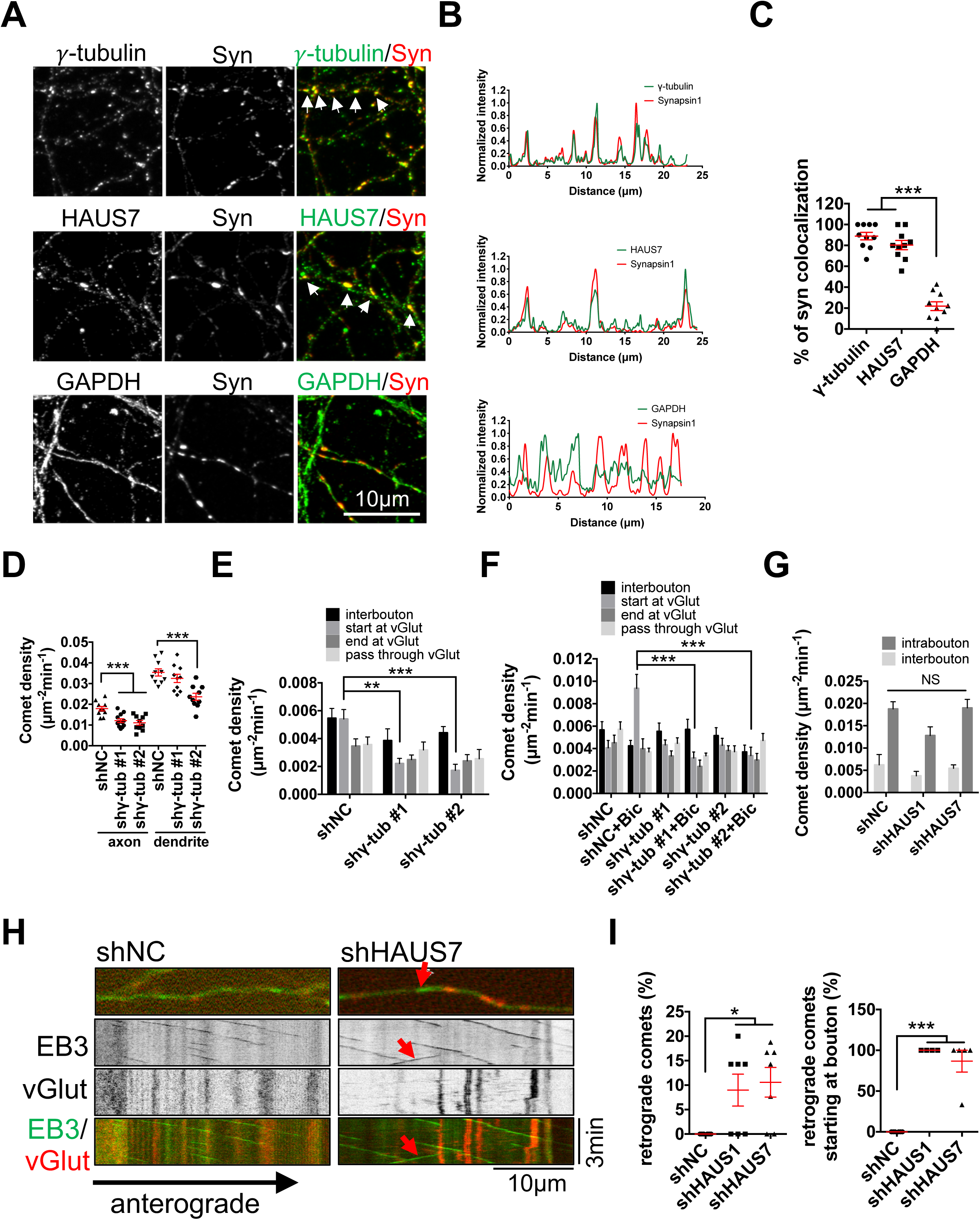
Neuronal activity induces *de novo* nucleation of distally oriented dynamic MTs through γ-tubulin and augmin function at presynaptic boutons. (A) Maximum projection of laser scanning confocal fluorescence images acquired with Airyscan detector in hippocampal neurons (21DIV) fixed and stained for γ-tubulin, HAUS7, GAPDH and synapsin1 (Syn). White arrows indicate signals co-localizing with Syn. (B) Normalized intensity of line scan of axons shown in (A) and indicated by arrows. (C) Quantification of percentage of Syn^+^ puncta co-localizing with γ-tubulin, HAUS7 or GAPDH. (D) Quantification of EB3 comet density in axons and dendrites of hippocampal neurons (20DIV) infected with noncoding control (shNC) or 2 different shRNAs against γ-tubulin (shγ-tub #1 or shγ-tub #2) for 6d. (E) Quantification of subclassified EB3 comet density relative to stable vGlut puncta in axons of hippocampal neurons as in (D). (F) Quantification of subclassified EB3 comet density relative to stable vGlut^+^ puncta of hippocampal neurons (18DIV) infected with shNC or shγ-tub#1 or #2 for 4d and transfected with EB3-EGFP (EB3) and vGlut1-mCherry (vGlut) 24h prior to live imaging. Neurons were pretreated with D-AP5 6-12h prior to imaging, followed by a washout and incubation with bicuculline for 1-10min (+Bic). (G) Quantification of intra- and interbouton comet density in neurons (21DIV) infected with shNC or shRNAs against HAUS1 or HAUS7 for 7d and transfected with EB3 and vGlut 24h prior to live imaging. (H) Kymographs of hippocampal neurons (21DIV) treated as in (G). Red arrows show a retrogradely oriented EB3 comet initiating from a bouton. (I) Quantification of the percentage of retrograde comets and retrograde comets starting at boutons of neurons as in (G). * p<0.05; ** p<0.01; *** p<0.001 by ANOVA (C: N = 10 axons; D-F: N = 10-12 axons) or one-tailed Student’s t-test (G, I: N=6-7 axons). NS, non significant.

When we knocked down endogenous γ-tubulin in hippocampal neurons for 6d with 2 different shRNAs, we achieved either ∼50% (shγ-tub #1) or ∼60% (shγ-tub #2) protein depletion and observed a significant decrease in comet density in both axons and dendrites (Figure 2D and Figure S2C-E). Interestingly, the decrease in comet density in axons was most significantly from loss of EB3 comets starting at presynaptic boutons (Figure 2E). Silencing of γ-tubulin for 4d, a condition that did not interfere with comet density or MT stability in unstimulated neurons (Figure 2F and Figure S2F,G), significantly abrogated the increase in the density of EB3 comets starting at boutons upon induction of neuronal activity (Figure 2F). These data strongly indicate that dynamic MTs initiating at excitatory boutons are a result of γ-tubulin dependent *de novo* nucleation and that presynaptic boutons are hotspots for activity-evoked MT nucleation in axons.

Nearly all newly nucleated MTs at presynaptic boutons displayed a biased growth orientation towards the distal end of the axon. The augmin complex regulates the polarity of γ-tubulin-nucleated MTs in nonneuronal cells and in axons of developing neurons, with γ-tubulin providing the nucleating material and the augmin complex being important for the maintenance of uniform polarity of axonal MTs (*24, 25*). Endogenous augmin complex subunit HAUS7 co-localized preferentially with synapsin1^+^ boutons compared to GAPDH control (∼80% versus ∼20%) (Figure 2A,B). Silencing of the augmin complex expression by knocking down subunits HAUS1 or HAUS7 to an extent (∼40% of protein depletion) that had no effect on comet density *per se,* significantly increased the percentage of retrograde comets, almost all of which initiated at boutons (Figure 2G-I, Figure S2H, and Video S4).

Collectively, these data indicate that γ-tubulin and augmin are preferentially localized at excitatory *en passant* boutons where they are required for activity-dependent *de novo* nucleation of uniformly distally-oriented dynamic MTs.

### MT nucleation at boutons is required for SV interbouton bidirectional motility and neurotransmission

We investigated whether *de novo* nucleated dynamic MTs at boutons could provide the tracks for activity-evoked motility of axonal cargos (*26–29*). To this end, we analyzed the dynamics of a wide range of axonal organelles as well as SVs relative to vGlut1^+^ stable presynaptic boutons in a time window during which we observed MT nucleation at boutons upon induction of neuronal activity. No substantial change was detected in Rab5 (early endosome marker; Figure S3A), Rab7 (late endosome marker; Figure S3B), LAMP1 (lysosome marker; Figure S3C), or mitochondria dynamics (Figure S3D), indicating that motility of these organelles is not immediately affected by activation of neuronal activity induced by bicuculline. We observed, however, a significant increase in the percentage of motile vGlut1^+^ and synaptophysin^+^ SVs that would start or end at stable presynaptic boutons (Figure 3A-B), with no preference in the direction of the movement (Figure S3E,F).

**Figure 3.**
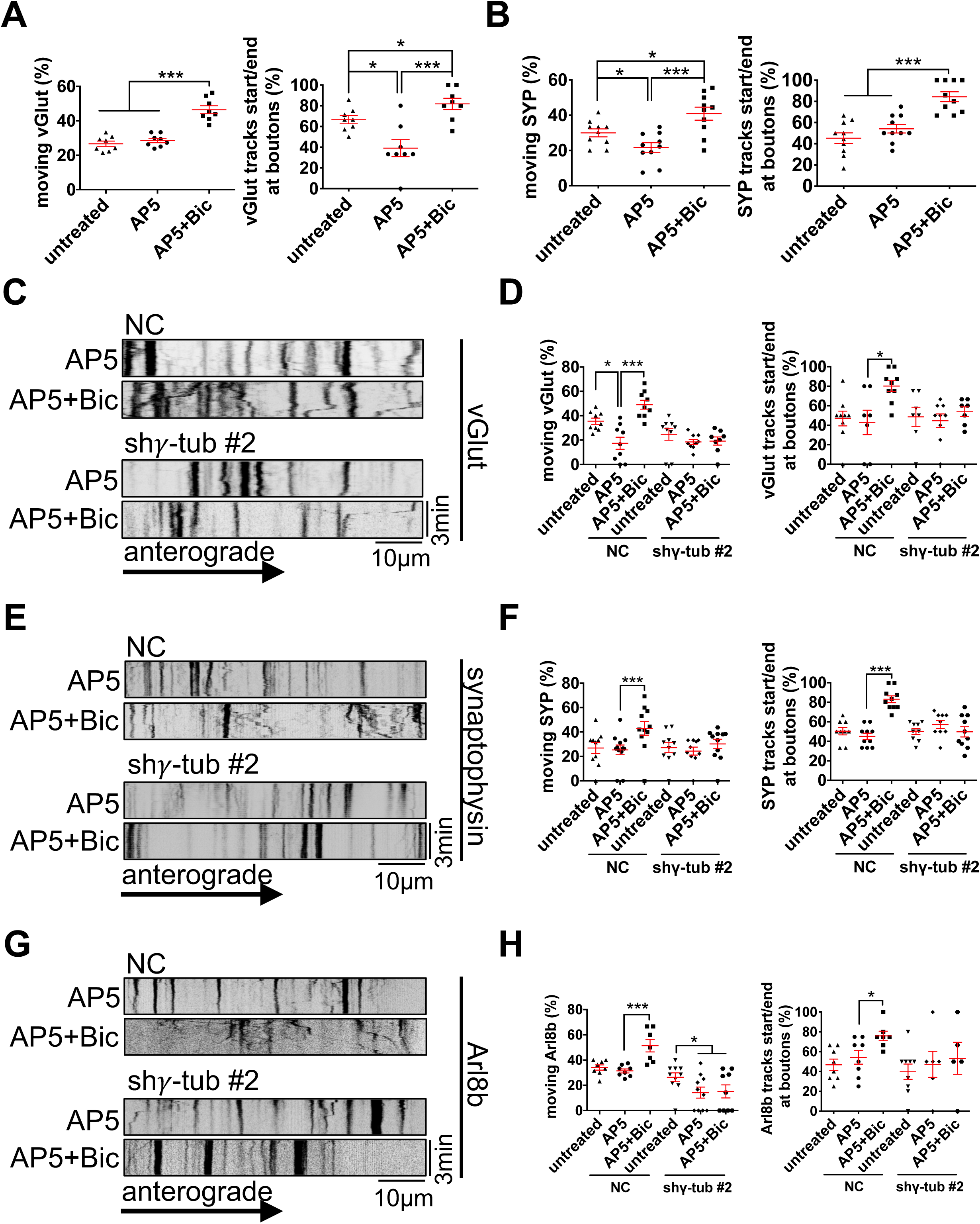
MT nucleation at boutons is required for bidirectional SV interbouton movement stimulated by neuronal activity. (A, B) Quantification of the percentage of moving vGlut^+^ (A) or synaptophysin^+^ (SYP) (B) puncta and percentage of moving tracks starting or ending at boutons in untreated hippocampal neurons (18DIV) or AP5 pretreated neurons followed by a washout with or without bicuculline. Neurons were transfected with EB3-EGFP and vGlut1-mCherry (A) or EB3-tdTomato and synaptophysin-Venus (SYP) (B), and vGlut1-mTAGBFP2 24h prior to live imaging. (C, E, G) Representative kymographs of vGlut (C), SYP (E), or Arl8b (G) in hippocampal neurons (18DIV) infected with noncoding control (shNC) or shRNA against γ-tubulin (shγ-tub #2) for 4d and treated as in A, B, or transfected with EB3-tdTomato and Arl8b-EGFP. (D, F, H) Quantification of the percentage of moving vGlut^+^ (D), SYP^+^ (F), or Arlb^+^ (H) puncta and the percentage of moving tracks starting or ending at boutons in hippocampal neurons treated as in C, E, or G. * p<0.05; ** p<0.01; *** p<0.001 by ANOVA (N = 8-12 axons).

To further understand whether local SV transport is dependent on *de novo* MT nucleation, we examined the mobile pool of vGlut1^+^, synaptophysin^+^ and Arl8b^+^ SVs in neurons depleted of γ-tubulin expression to levels that would only affect activity evoked MT nucleation at boutons. Remarkably, we found that γ-tubulin knockdown strongly inhibited the activity evoked motility of SVs towards and away from the boutons (Figure 3C-H). This demonstrates that activity evoked MT nucleation at boutons is required for stimulated SV targeted interbouton delivery.

Inhibition of Arl8-mediated transport of SVs and presynaptic lysosome related vesicles (PLVs) affects bouton size and neurotransmission during bouton biogenesis (*30, 31*). Thus, we examined whether dynamic MTs nucleated at boutons could affect SV pool size and/or exocytosis kinetics at presynaptic release sites of neurons with mature synapses. To this end, the pH sensitive vesicular chimeric marker vGlut1-pHluorin and the cytosolic filler mCherry-C1 were co-transfected into neurons (16DIV) that had been depleted of γ-tubulin expression by two independent shRNAs to levels that would only affect evoked-MT nucleation at boutons. We measured no difference in vGlut1-pHluorin signal intensity at active presynapses after alkalinization with NH4Cl. However, a marked reduction of stimulated fluorescence in γ-tubulin deprived neurons was observed compared to control neurons, indicating that loss of AP-evoked MT nucleation at boutons strongly inhibited stimulated exocytosis without altering the total pool of SVs residing at boutons (Figure 4A-E).

**Figure 4.**
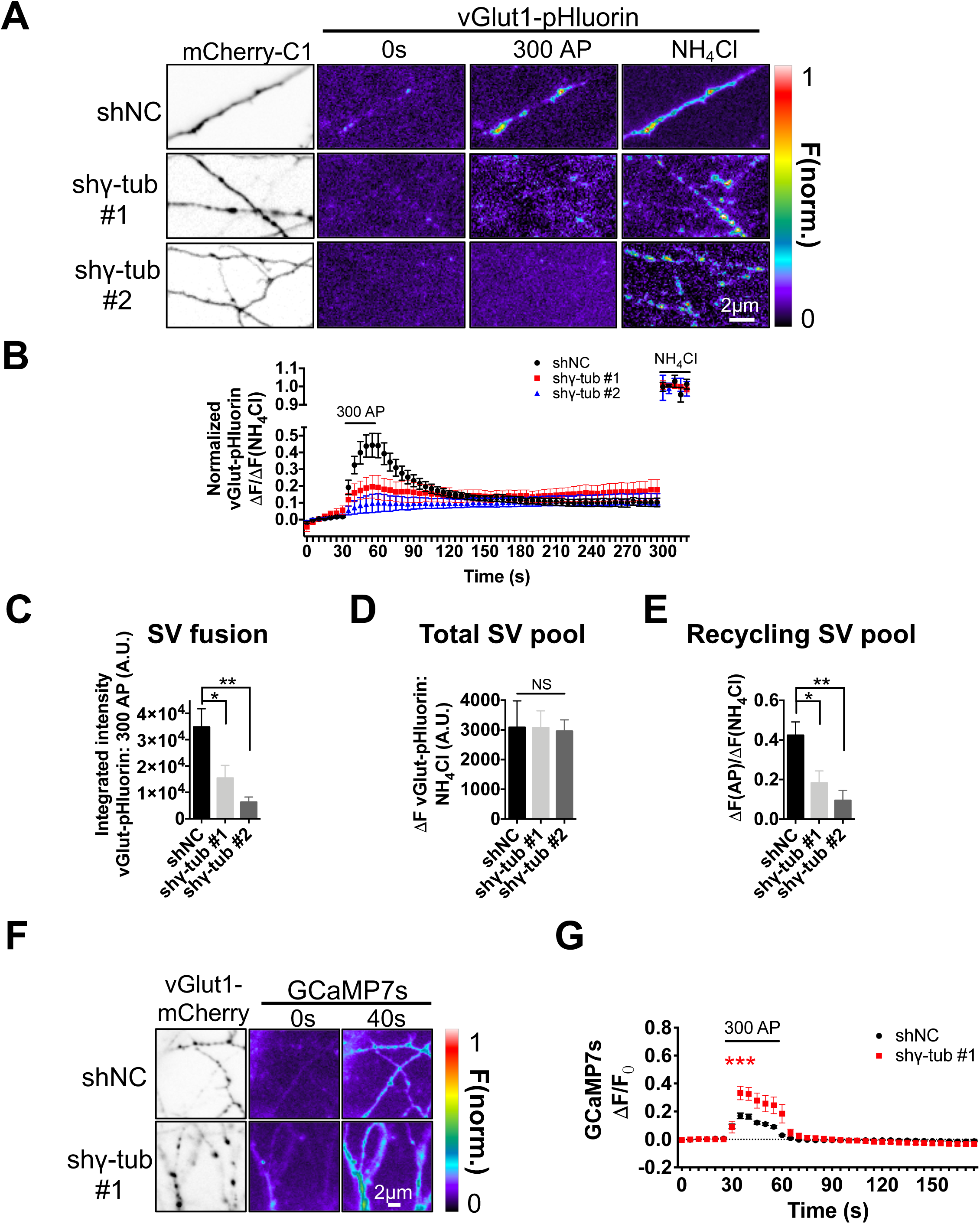
MT nucleation at boutons is required for SV exocytosis. (A). Representative images of hippocampal neurons (16DIV) infected with noncoding control (shNC) or two independent shRNAs against γ-tubulin (shγ-tub #1 or #2) for 4d and co-transfected with vGlut1-pHluorin and mCherry-C1 prior to recording vGlut1-pHluorin dequenching for the indicated times in response to neuronal activation induced by 300 action potentials (AP) at 10Hz starting from 30s. (B) Normalized fluorescence change (δF) over total vGlut1-pHluorin (δF upon NH4Cl dequenching) at the indicated times. (C) Integrated intensity of vGlut1-pHluorin signal during the 300AP stimulation paradigm. (D) Total pool of SVs indicated by vGlut1-pHluorin signal intensity at active presynapses after alkalinization with NH_4_Cl. (E) Recycling pool of SVs indicated by the ratio of δF at maximum stimulation over δF after alkalinization with NH_4_Cl. (F) Representative images of hippocampal neurons (16DIV) infected shNC or shγ-tub #2 lentiviruses for 4d and co-transfected with GCaMP7s and vGlut1-mCherry using the same stimulation protocol as in A. (G) Ratio of GCaMP fluorescence change (Δ F) over baseline intensity (Δ F) at the indicated times. * p<0.05; ** p<0.01; *** p<0.001 by two-tailed Student’s t-test (A-E: N = 4-6 imaging fields including 8-18 axons with 200-360 boutons; F,G: N = 3-4 imaging fields including 6-12 axons with 120-150 boutons). Red asterisk in G compares γ-tubulin #2 to control levels at 40s.

Ca^2+^ influx through voltage-gated Ca^2+^ channels plays a crucial role in exocytosis, and the strong inhibition of AP-triggered exocytosis upon γ-tubulin depletion could potentially be due to a decreased Ca^2+^ entry through voltage-gated Ca^2+^ channels. We examined the impact of loss of γ-tubulin expression on AP-evoked Ca^2+^ influx (*32*), and found that depleting γ-tubulin expression for 4d did not inhibit AP-evoked Ca^2+^ influx, ruling out suppression of Ca^2+^ entry as a mechanism of the observed inhibition of presynaptic release (Figure 4F,G).

Together, these data indicate that activity stimulated *de novo* MT nucleation at boutons is required for SV interbouton transport and neurotransmitter release.

## DISCUSSION

We found that in primary hippocampal neurons, excitatory *en passant* boutons are hotspots for γ-tubulin dependent MT nucleation that is induced by neuronal activity. Importantly, MT nucleation preferentially occurs at excitatory boutons in hippocampal slices from neonatal mice, demonstrating that localized MT nucleation at boutons is *a bona fide* feature of mature hippocampal neurons residing in tissue where cytoarchitecture and synaptic organization are preserved (Figure 1).

γ-tubulin had been shown to control acentrosomal MT nucleation, neuronal polarity and axonal outgrowth in developing dendrites and axons (*25, 33*). The present study reports a nucleating function of γ-tubulin in neurons with mature synapses that is compartmentalized, regulated by neuronal activity and required for synaptic transmission.

Acute γ-tubulin knockdown only affected axonal MT nucleation from the boutons, with negligible effects on interbouton comet density or catastrophe frequency (Figure 2D-F and Figure S2C-E). Although this may reflect incomplete γ-tubulin depletion even after 6d of silencing, our data suggest that γ-tubulin is specifically required for MT nucleation at the boutons in response to neuronal activity and that interbouton comet density is regulated by either rescue factors or a yet unknown non-γ-tubulin dependent MT nucleating machinery such as the recently discovered MT remodeling factor SSNA1 (*34*). Furthermore, given the evidence that ∼80% of γ-tubulin^+^ puncta are localized at *en passant* presynaptic boutons, we speculate that not all boutons are selected to nucleate MTs, and that this function depends on regulated γ-tubulin localization and/or activation. The basis for this selectivity is at moment unknown but may reflect modulation of SV releasing activity at individual synapses, and/or the ability of a bouton to establish functional contacts with a postsynaptic site. Future investigation is needed to detail the mechanisms of γ-tubulin recruitment to presynaptic boutons and the rules establishing bouton selection.

We further demonstrate that the correct polarity of presynaptic *de novo* nucleated MTs requires the augmin complex and that similarly to γ-tubulin ∼80% of augmin^+^ puncta are localized at *en passant* presynaptic boutons. Interestingly, augmin complex depletion had no impact on MT polarity in dendrites as antiparallel MT organization could be regulated by other mechanisms (*2, 24*). Indeed, the augmin complex has been shown to generate local MTs with conserved polarity in developing axons (*24, 25*). Our results not only demonstrate that the augmin complex restricts the uniform orientation of dynamic MTs at presynaptic boutons of mature axons but further support the notion that these newly generated presynaptic MTs result from *de novo* γ-tubulin nucleation rather than MT rescue or SSNA1-mediated protofilament splitting. How mature axons and boutons preferentially recruit and/or activate the augmin complex in coordination with γ-tubulin for the regulation of MT nucleation at selected boutons upon neuronal firing remains to be established.

We found that activity-dependent MT nucleation at boutons is required for interbouton SV motility, and to allow neurotransmitter release (Figures 3 and 4). Arl8b facilitates the Kif1A mediated motility of both SVs and PLVs, which coordinate the anterograde transport of packets of AZ and SV proteins in developing neurons during presynaptic biogenesis (*30, 31, 35*). In addition, Kif1A was recently implicated in synaptic cargo anterograde delivery to *en passant* boutons using dynamic MTs as tracks (*23*). Based on this collective evidence, we propose a model in which dynamic MTs are *de novo* nucleated to serve as tracks for targeted interbouton delivery of a rate-limiting supply of SV and AZ precursors at sites of stimulated release. Indeed, our observation that neuronal activity stimulates presynaptic *de novo* nucleation of dynamic MTs that terminate to the next distal bouton is in line with the findings that *en passant* boutons are enriched in dynamic MT plus ends (*23*). Our results differ in that we observed that: (1) only MT nucleation (initiation) but not catastrophe (termination) is preferentially regulated at boutons and (2) both anterograde and retrograde SV transport are dependent on MT nucleation at boutons. The discrepancy may be explained because, in contrast to their measurements which spanned along the entire length of the axon, our *in vitro* and *ex vivo* comet tracking was restricted to proximal axons, a region where interbouton bidirectional transport of pools of SVs may be predominant (*23, 36–39*). The model that Guedes-Dias et al. propose may be more consistent with targeted long-distance delivery and retention of SV precursors to synapses residing in the most distal region of the axon. At proximal synaptic sites, however, a higher proportion of bidirectional long-distance delivery of SVs may be required in response to neuronal activity. This is consistent with our observation that in proximal axons ∼40% of nucleated comets at a bouton reach to the next bouton after bicuculline-induced firing compared to ∼10% before stimulation in dissociated hippocampal neurons (Figure S1C) or ∼25% in hippocampal slices.

We found that loss of *de novo* nucleated MTs by γ-tubulin depletion significantly impairs neurotransmitter release, paving the way for the characterization of the mechanisms by which presynaptic dynamic MTs may accomplish this task through targeted delivery of a rate-limiting pool of SVs to the sites of fusion and/or by mediating the transport of a rate-limiting vesicular component of the fusion or AZ machinery.

At present, the signaling pathways that regulate presynaptic MT nucleation are unknown. One hypothesis is that neuronal firing may drive Ca^2+^ influx dependent recruitment of MT nucleation complexes only at active boutons. Given the critical role that loss of a functional presynaptically-localized machinery for MT nucleation may have for neurotransmission, it will be important to determine the *in vivo* significance of this selective MT nucleation and whether dysregulation of this process can be associated with human neurological and neuropsychiatric illnesses caused by mutations in MT regulatory proteins residing presynaptically, such as tau in AD (*40, 41*) and tauopathies, spastin in hereditary spastic paraplegias (*42, 43*), and MAP1B and FMRP in Fragile X Syndrome (*44–46*).

## ACKNOWLEDGMENTS

Our gratitude goes to Franck Polleux for providing us with the *ex vivo* experimental set up, EB3-EGFP, mito-DsRed2, synaptophysin-Venus, and vGlut1-mTagBFP2 constructs in addition to helpful discussions; Viktoriya Zhuravleva for assisting us with the vGlut1-pHluorin experimental set up; Inbal Israely for sharing pCMV-GCaMP7s construct; Richard Vallee for sharing EGFP-Rab7a construct; Volker Haucke for sharing LAMP1-EGFP and ARl8b-EGFP constructs; Ryan Bose-Roy for helping with the analysis of the movies. We are grateful to Gregg G. Gundersen and Richard Vallee for access to their microscopes. This project was funded by an RO1AG050658 (NIH/NIA) to F.B..

## AUTHOR CONTRIBUTIONS

X.Q. and F.B. designed the project, analyzed all the data and wrote the manuscript. X.Q. performed most of the experiments. H.B. carried out the *in utero* hippocampal electroporation. A.K. and H.B. performed the *ex vivo* time-lapse imaging experiment with acute hippocampal slices. vGlut1-Phluorin and GCaMP7s experiments in hippocampal neurons were carried out by A.K. under C.W. supervision.

## SUPPLEMENTAL INFORMATION

**Figure S1.**
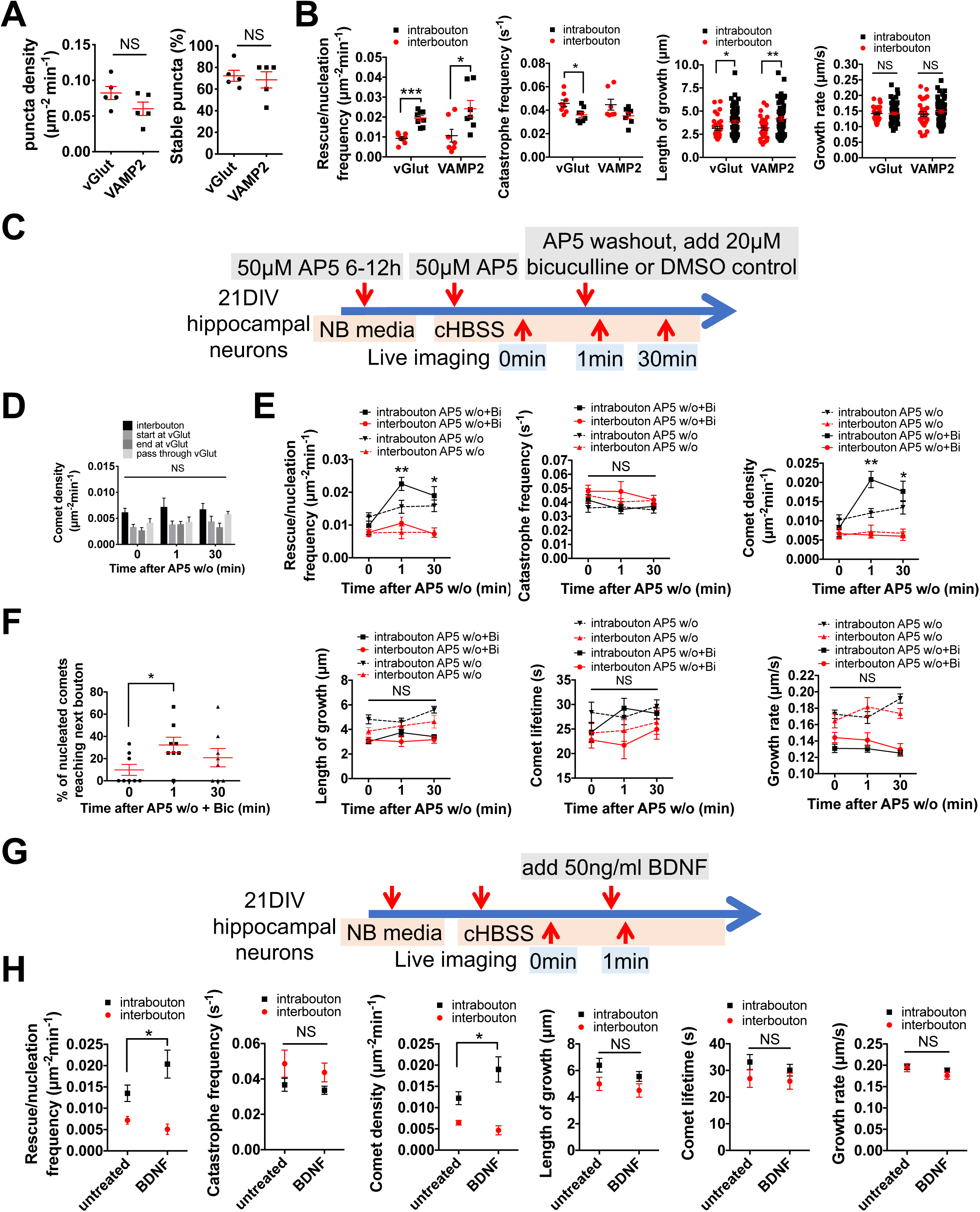
Additional MT dynamics parameters measured in untreated or treated hippocampal neurons (21DIV) transfected with EB3-EGFP and vGlut1-mCherry or VAMP2-Cherry. (A) Total density and percentage of stable vGlut1 and VAMP2 puncta in untreated hippocampal neurons (21DIV) transfected with EB3-EGFP and vGlut1-mCherry or VAMP2-Cherry. (B) Rescue/nucleation frequency, catastrophe frequency, length of growth, and growth rate of intrabouton or interbouton EB3 comets in movies of neurons as in A. (C) Schematics showing the timeline of neuronal activation using AP5 and bicuculline. (D) Subclassified comet density for intrabouton MTs measured in hippocampal neurons (21DIV) transfected with EB3-EGFP and vGlut1-mCherry and treated as in (C) using DMSO control. (E) Rescue/nucleation frequency, catastrophe frequency, comet density, length of growth, comet lifetime, and growth rate of intrabouton or interbouton EB3 comets in neurons treated as in (C) using bicuculline. (F) Percentage of comets nucleated at boutons that reach the next distal bouton in neurons treated as in (C) using bicuculline. (G) Schematics showing the timeline of neuronal activation by treating with 50ng/mL BDNF for 1min. (H) Rescue/nucleation frequency, catastrophe frequency, comet density, length of growth, comet lifetime, and growth rate of intrabouton or interbouton EB3 comets in neurons treated as in (G). * p<0.05; ** p<0.01; *** p<0.001 by two-tailed Student’s t-test (A, B: N = 5-8 axons; D, E, F, H: N = 6-9 axons). NS, non significant.

**Figure S2.**
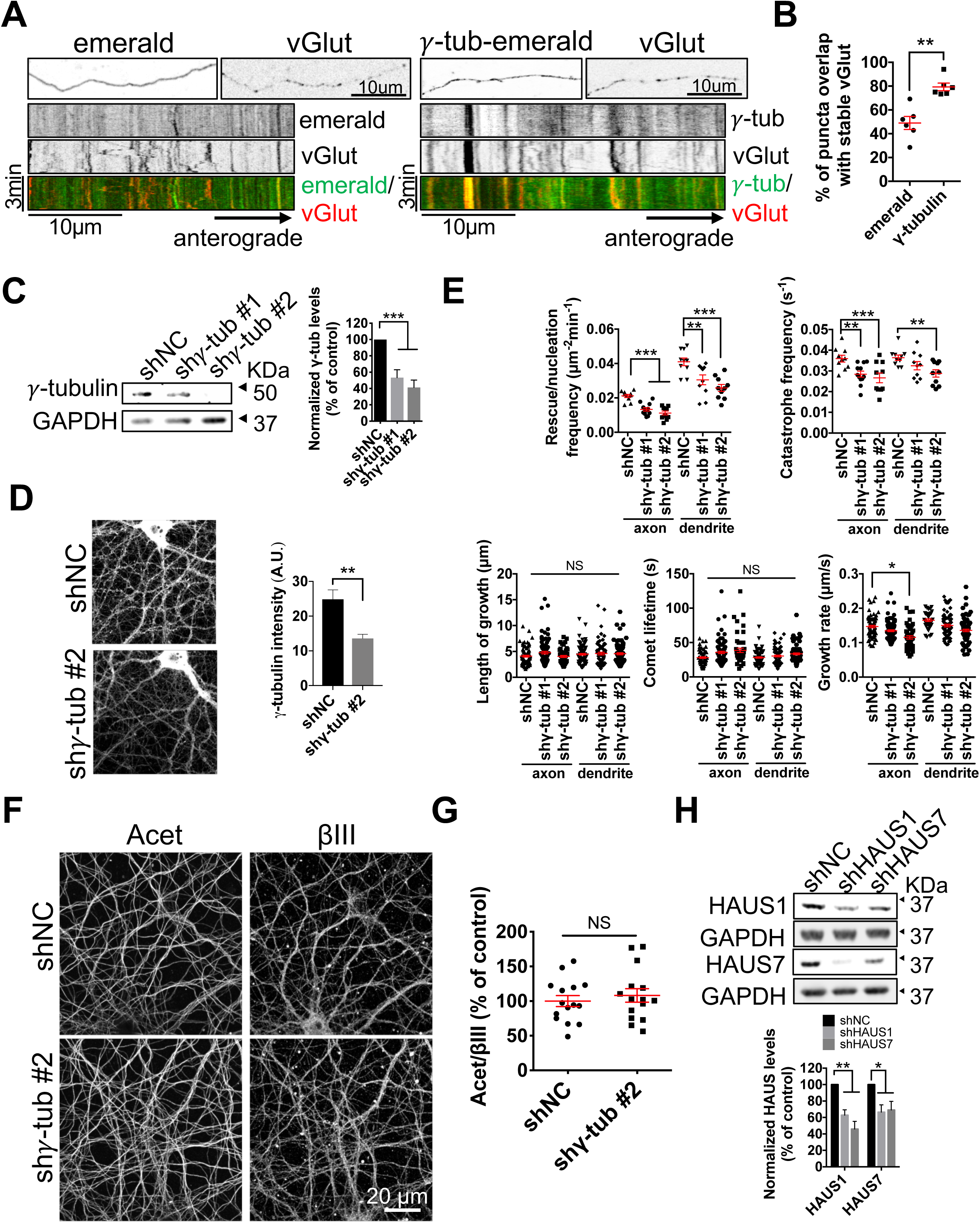
Additional co-localization of exogenous expressed γ-tubulin and vGlut1 and analysis in γ-tubulin or HAUS knocked down hippocampal neurons. (A) Maximum projection of spinning disk confocal fluorescence images and kymographs of hippocampal neurons (21DIV) transfected with emerald control or γ-tubulin-emerald and vGlut-mCherry (vGlut). (B) Quantification of the percentage of emerald control or γ-tubulin-emerald signal co-localizing with stable vGlut-mCherry (vGlut) puncta in neurons as in (A). (C) Western blot analysis of γ-tubulin levels in hippocampal neurons (20DIV) infected with noncoding control (shNC) or 2 different shRNAs against γ-tubulin (shγ-tub #1 or shγ-tub #2) for 6d. (D) Quantitative immunofluorescence of γ-tubulin ackground subtracted intensity in hippocampal neurons (20DIV) infected with shNC and shγ-tub #2 for 6d. (E) MT dynamics analysis of dendrites and axons in hippocampal neurons treated as in (C). (F) Immunofluorescence of acetylated (Acet) and βIII tubulin in hippocampal neurons (18DIV) infected with noncoding control (shNC) or shRNAs against γ-tubulin (shγ-tub #2) for 4d. (G) Ratio analysis of normalized Acet relative to βIII tubulin intensity. (H) Western blot analysis of HAUS1 and HAUS7 levels in hippocampal neurons (21DIV) infected with shNC or shRNAs against HAUS1 or HAUS7 for 7d. * p<0.05; ** p<0.01; *** p<0.001 by two-tailed Student’s t-test (B: N = 6 neurites; C: N = 5 experiments; D: N = 4 fields of view; E: N = 10-12 neurites; G: N = 15 axons; H: N = 3 experiments). NS, non significant.

**Figure S3.**
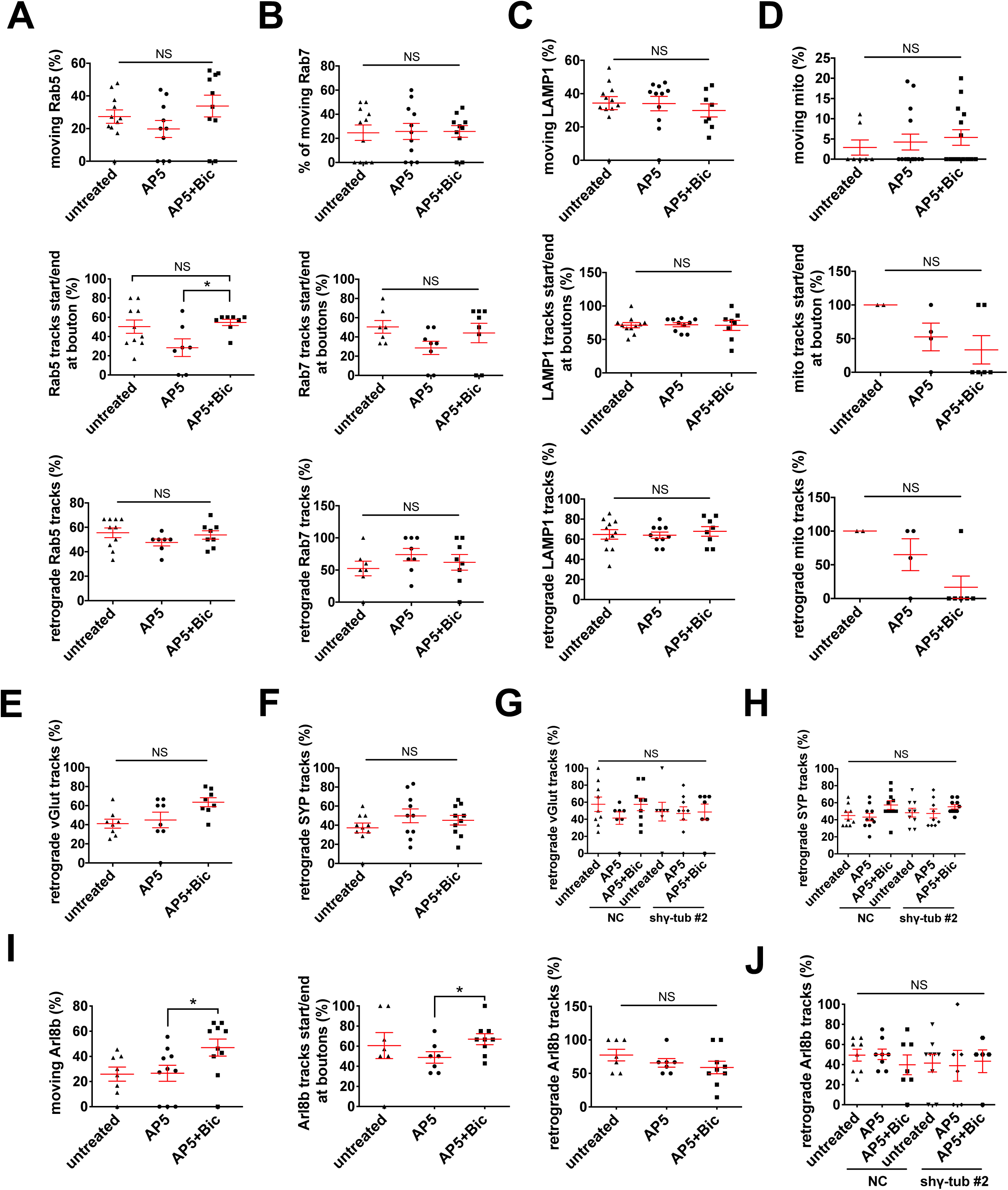
Selected axonal organelle dynamics are mostly unaffected by acute stimulation of neuronal activity. (A, B, C, D) Quantification of the percentage of moving, retrograde or bidirectional moving tracks starting or ending at boutons of Rab5 (A), Rab7 (B), LAMP1 (C), and mitochondria (D) in hippocampal neurons transfected with each organelle marker, EB3-EGFP/EB3-tdTomato and vGlut-mTAGBFP2 in shNC or shγ-tub #2 infected hippocampal neurons (18DIV) for 4d prior to imaging under untreated condition, or pretreated with AP5 for 6h followed by a washout with or without bicuculline. (E, F, G, H, J) Percentage of retrograde vGlut (E, G), SYP (F, H), Arl8b (J) tracks in non-infected (E, F), shNC or shγ-tub #2 infected (G, H, J) hippocampal neurons transfected with vGlut1-mCherry/synaptophysin-venus/Arl8b-EGFP, EB3-EGFP/EB3-tdTomato and vGlut-mTAGBFP2, and treated as in (A). (I) Quantification of the percentage of moving, retrograde or bidirectional moving tracks starting or ending at boutons of Arl8b in hippocampal neurons transfected with Arl8b-EGFP, EB3-Tdtomato, and vGlut-mTAGBFP2 and treated as in (A). * p<0.05 by two-tailed Student’s t-test (N > 7 axons). NS, non significant.

**Video S1**

Maximum intensity projection of a time-lapse showing EB3-EGFP comet motility relative to stable vGlut1-mCherry labeled boutons from an axon residing in the CA1 hippocampal region of an acute slice.

**Video S2**

Maximum intensity projection of a time-lapse showing EB3-EGFP comet motility relative to stable vGlut1-mCherry labeled boutons from an axon of a cultured hippocampal neuron pretreated with 50µM AP5 followed by a washout and incubation with 20µM bicuculline.

**Video S3**

Maximum intensity projection of a time-lapse showing EB3-EGFP comet motility relative to stable vGlut1-mCherry labeled boutons from an axon of a cultured hippocampal neuron treated with 50ng/ml BDNF.

**Video S4**

Maximum intensity projection of a time-lapse showing EB3-EGFP comet motility relative to stable vGlut1-mCherry labeled boutons from axons of cultured hippocampal neurons infected with shNC control or shHAUS7 lentiviruses for 7d.

## EXPERIMENTAL PROCEDURES

### Animals

All protocols and procedures were approved by the Committee on the Ethics of Animal Experiments of Columbia University and according to Guide for the Care and Use of Laboratory Animals of the National Institutes of Health. E18 pregnant Sprague Dawley rats were purchased from Charles River Laboratories.

### Reagents and antibodies

All chemicals were obtained from Sigma-Aldrich (St. Louis, MO), unless otherwise noted. Primary antibodies used for western blotting (WB) or immunofluorescence (IF) were: mouse anti-γ tubulin (TU-30, Exbio, 1:1000 WB, 1:100 IF), mouse anti-GAPDH (1:5000 WB), rabbit anti-GAPDH (1:5000 WB, 1:100 IF), rabbit anti-HAUS1 (ThermoFisher, 1:1000 WB), mouse anti-HAUS7 (OTI1E8, ThermoFisher, 1:1000 WB, 1:100 IF), mouse anti-acetylated tubulin (6-11-B1, 1:100 IF), rabbit anti-βIII-tubulin (Abcam, 1:1000 IF), Rabbit anti-synapsin1 (SYSY, 1:1000 IF), mouse anti-synapsin1 (SYSY, 1:1000 IF). For western blot analysis, secondary antibodies were conjugated to IR680 or IR800 (Rockland Immunochemicals) for multiple infrared detection. For immunofluorescence analysis, primary antibodies incubation was followed by use of the appropriate Alexa Fluor fluorescent dyes-conjugated secondary antibodies (Jackson Immunoresearch Laboratories).

### Primary hippocampal neuronal cultures

Primary hippocampal neuronal cultures were prepared as previously described (*47*). Briefly, hippocampi were dissected from E18 rat, and neurons plated on 100 μg/mL poly-D-lysine-coated 12-well-plates at the density of 3 × 10^5^ cells/well for biochemistry assays, 5 × 10^4^ cells/dish for live imaging in the chamber of 35mm MatTek dishes, or 4 × 10^4^ cells/coverslip on18 mm coverslips for immunofluorescence. Primary neurons were maintained in Neurobasal medium (Invitrogen) with the supplement of 2% B-27 (Invitrogen) and 0.5mM glutamine (Invitrogen), and 1/3 of the medium was changed every 3-4d up to 3w in culture.

### Lentiviral shRNA silencing

Production of lentiviral particles was conducted using the 2^nd^ generation packaging system as previously described (*47*). Briefly, HEK293T were co-transfected with lentiviral shRNA constructs and the packaging vectors pLP1, pLP2, and pLP-VSV-G (Invitrogen) using calcium phosphate. 48h and 72h after transfection, the virus was collected, filtered through 0.22μm filter, and further concentrated by ultracentrifugation. Concentrated virus was aliquoted and stored at −80 °C. Lentiviral constructs to knockdown γ-tubulin were purchased from Sigma Aldrich (TRCN0000089905 and TRCN0000089907) with the following DNA sequence 5’-CCGGGCAATCAGATTGGGTTCGAGTCTCGAGACTCGAACCCAATCTGATTGCTTTTTG-3’ and 5’-CCGGGCAGCAGCTGATTGACGAGTACTCGAGTACTCGTCAATCAGCTGCTGCTTTTTG-3’ onto pLKO.1 lentivector. Lentiviral constructs to knockdown HAUS1 and HAUS7 were purchased from Sigma Aldrich (TRCN0000192324 and TRCN0000345686) with the following DNA sequence 5’-CCGGCTTTCTCATGGAGAGTGTGAACTCGAGTTCACACTCTCCATGAGAAAGTTTT TTG-3’ and 5’-CCGGCCAGATGACCAGGATCTTCTACTCGAGTAGAAGATCCTGGTCATCTGGTTTTTG-3’ onto pLKO.1 lentivector. The pLKO.1 vector with noncoding (NC) sequence was used as control.

### Western blot analyses

Cells were lysed in Laemmli sample buffer and boiled at 96°C for 5m. Cell lysates were sonicated by a probe sonicator to sheer cellular debris and genomic DNA. To detect tubulins, lysates were diluted 1:10 in sample buffer. Proteins were separated by 10% Bis-Tris gel (Invitrogen) and transferred onto nitrocellulose membrane. After blocking in 5% milk/TBS or BSA/TBS, membranes were incubated with primary antibodies at 4°C overnight prior to 1h incubation with secondary antibodies. Image acquisition was performed with an Odyssey imaging system (LI-COR Biosciences, NE) and analyzed with Odyssey software.

### Immunofluorescence microscopy and analyses

For MT staining, neurons were fixed in 3.7% PFA, 0.25% glutaraldehyde, 3.7% sucrose and 0.1% triton-X 100 for 15 min and quenched in 1 mg/mL sodium borohydride/PBS. For γ-tubulin, HAUS7, GAPDH and synapsin staining, neurons were fixed in 4% PFA for 15 min. Cells were then washed in PBS, blocked in 2% FBS, 2% BSA, and 0.2% fish gelatin in PBS for 1 h, and stained with primary antibodies for 2h followed by secondary antibodies for 1h. Mounted samples were observed by a Zeiss LSM 800 confocal microscope equipped with Airyscan module, using a 63x objective (Plan-Apochromat, NA 1.4). Images were obtained and processed for super resolution using Zen Blue 2.1 software. All images were analyzed by ImageJ software except for pseudocolor presentation of vGlut-pHluorin experiments that were instead obtained using Metamorph software.

### Live imaging of MT, organelle and SV dynamics

Neurons grown on MatTek dishes were co-transfected with EB3-EGFP or EB3-tdTomato using Lipofectamine 2000 (Invitrogen), together with presynaptic markers vGlut1-mCherry, vGlut1-Venus, vGlut1-mTagBFP2, or VAMP2-mCherry. For organelle dynamics, triple transfection of EB3, vGlut1, together with the organelle markers Rab5-EGFP, EGFP-Rab7a, LAMP1-GFP, or mitoDsRed were performed using Lipofectamine 2000. Live cell imaging was performed 24-48h after transfection in complete HBSS media (HBSS, 30mM glucose, 1mM CaCl_2_, 1mM MgSO_4_, 4mM NaHCO_3_, and 2.5mM HEPES, pH 7.4) using IX83 Andor Revolution XD Spinning Disk Confocal System. The microscope was equipped with a 100×/1.49 oil UApo objective, a multi-axis stage controller (ASI MS-2000), and a controlled temperature and CO_2_ incubator. Movies were acquired with an Andor iXon Ultra EMCCD camera and Andor iQ 3.6.2 live cell imaging software at 0.5-2 s/frame for 3min. For pharmacological induction of neuronal activity, neurons were pretreated with 50μM D-AP5 for 6-12h prior to live imaging in complete neurobasal media, then changed to complete HBSS media with the same concentration of D-AP5 for live imaging. To induce neuronal activity, neurons were washed 3x with complete HBSS media and 20μM bicuculline or DMSO control was added to complete HBSS media after washes. For BDNF induction of neuronal activity, 50ng/ml BDNF was directly added to complete HBSS media during live imaging. Movies were taken 1min upon treatment. Maximum projections of movies were performed by Image Math within Andor software, exported as Tiff files and analyzed in ImageJ.

Kymographs were generated by drawing a region from the proximal to the distal axons (within 100μm from the cell body) based on morphology and anterograde movement of EB3-labeled comets. Presynaptic MTs were classified based on their plus end contacts with stable vGlut1 or VAMP2 labeled boutons. Parameters describing MT dynamics were defined as follows: rescue/nucleation frequency: number of rescue or nucleation events per μm^2^ per min; catastrophe frequency: number of full tracks/total duration of growth; comet density: number of comets per μm^2^ per min; growth length: comet movement length in μm; comet lifetime: duration of growth; growth rate: growth length/comet lifetime (*48*). Parameters describing organelle and SV dynamics were defined as follows: % of moving puncta: number of moving puncta/number of total puncta × 100; % of tracks start/end at boutons: number of tracks start or end at boutons/number of total moving tracks × 100; % of retrograde tracks: number of retrograde tracks/number of total moving tracks × 100.

### *In utero* electroporation and live-imaging of acute hippocampal slices

A mix of endotoxin-free plasmid EB3-EGFP (1.5mg/mL) and vGlut-mcherry (1.2mg/mL) and 0.5% Fast Green (Sigma) was injected into one lateral hemisphere of E15.5 embryos using a Picospritzer III (Parker). Electroporation (ECM 830, BTX) was performed with 5mm electrode tweezers (Nepagene, Japan) to target hippocampal progenitors in E15.5 embryos using a triple electrode setup according to dal Maschio et al. (*49*). Five pulses of 45V for 50ms with 500ms interval were used for electroporation. Animals were sacrificed 21-28 days after birth. Acute hippocampal slices (200µm) were prepared using a vibratome and collected in complete HBSS media (HBSS, 30mM glucose, 1mM _CaCl2,_ 1mM MgSO_4_, 4mM NaHCO_3_, and 2.5mM HEPES, pH 7.4). Acute slices were immediately transferred in 35mm MatTek dishes for live imaging. Electroporated hippocampal neurons from CA1 region were selected and imaged using an inverted Nikon Ti-E microscope with Nikon Elements Software at 37°C. 488 and 561nm lasers were used as light source and shuttered by Acousto-Optic Tunable Filters (AOTF). Live-imaging acquisition of EB3 comets relative to vGlut1 positive stable puncta within 50-100 µm of cell bodies was performed for 300s using a 60x oil-immersion objective (NA1.4) at 3 s/frame.

### Live imaging of SV exo-/endocytosis

To test synaptic exo-endocytosis via the vGlut-pHluorin assay using electrical field stimulation, hippocampal neurons were infected with noncoding shNC control or two independent shRNAs against γ-tubulin (shγ-tub #1 or #2) lentiviruses at 12DIV and co-transfected with vGlut-pHluorin and mCherry at 13DIV. Imaging of live neurons was performed at 16DIV in Tyrode’s buffer containing 119mM NaCl, 2.5mM KCl, 2mM MgCl2, 2mM CaCl2, 25mM Hepes, pH 7.4, 30mM D-glucose, 10µM 6-cyano-7-nitroquinoxaline-2,3-dione (CNQX; Tocris), and 50µM D-(-)-2-amino-5-phosphonopentanoic (D-AP5 or APV; Tocris) buffered to pH 7.4 at 37°C. Images were acquired with a 40X objective (Neofluar, NA 1.3) on an epifluorescence microscope (Axio Observer Z1, Zeiss) with Colibri LED light source, EMCCD camera (Hamamatsu) and Zeiss Zen software. Stills of the mCherry and pHluorin channels were acquired followed by a 300s time lapse of the pHluorin channel recorded at 5s/frame. Action potentials (AP) were evoked by passing 1msec bipolar current through platinum-iridium electrodes connected to metal tubes containing imaging buffer at 10Hz using a digital stimulator (World Precision Instruments) and a stimulator isolator (World Precision Instruments). Tyrode’s buffer supplemented with 50mM NH4Cl was perfused at the end of the recording to assess maximum pHluorin signal.

### Live imaging of Ca^2+^ influx by GCaMP7s

Primary hippocampal neurons (16DIV) were infected with noncoding shNC control or two independent shRNAs against γ-tubulin (shγ-tub #1 or #2) lentiviruses at 12DIV and co-transfected with GCaMP7s and vGlut1-mCherry at 13DIV. Live imaging was performed at 16DIV using the same microscopy setup and protocol as SV exo-/endocytosis without NH4Cl dequenching at the end of each recording.

### Statistical analyses

Data are shown as means ± SEMs from at least 3 independent experiments. Statistical analysis between two groups was performed using Student’s t tests. Comparison among 3 or more groups was performed using one-way ANOVA with Tukey’s post-test or Dunnett’s multiple comparisons test. Statistical significance was set for p<0.05.

